# Detection and Characterization of West Nile Virus with evidence of Transovarial Transmission in the Coastal region, Kenya

**DOI:** 10.64898/2025.12.01.691525

**Authors:** Tabitha Wanjiru, Solomon Langat, Hellen Koka, Santos Yalwala, Gladys Kerich, Janet Ambale, Jaree Johnson, Eric Garges, Robert Haynes, Gerald Kellar, John Eads, Fredrick Eyase

## Abstract

**Background:** West Nile virus (WNV) is a mosquito-borne flavivirus of global public health significance, maintained in an enzootic cycle between birds and mosquitoes, with humans and other mammals as incidental hosts. Understanding WNV circulation in diverse mosquito populations is critical for predicting and mitigating outbreaks. This study investigated mosquito populations from coastal Kenya, where WNV was detected, genetically characterized, and evidence of transovarial transmission in Aedes aegypti was observed.

**Methods:** Mosquitoes were collected from Kwale, Kilifi, Mombasa, and Isiolo counties (n=14,105) and pooled by species and location (1,596 pools). Pools were inoculated on Vero E6 cells, followed by RNA extraction, Illumina MiSeq sequencing, and preliminary analysis on CZ-ID. Reads were quality-controlled (PrinseqLite v0.20.4), assembled de novo (MEGAHIT v1.2.9), and analyzed via BLAST. Phylogenetic reconstruction used Maximum Likelihood, and codon-level selection pressure was evaluated using FEL, MEME, and FUBAR on Datamonkey.

**Results:** WNV was detected in ten pools: eight Lineage 1a and two Lineage 2. Virus isolates came from *Culex pipiens*, *Culex univittatus*, *Anopheles funestus*, *Aedes aegypti*, and *Eretmapodites chrysogaster*. Notably, one Lineage 1a isolate from a male *Aedes aegypti* confirmed transovarial transmission. Six codons were under diversifying selection, NS2B gene was found to carry the V103A mutation.

**Conclusion:** WNV in coastal Kenyan mosquitoes revealed transovarial transmission in *Aedes aegypti*. Lineage-specific evolution, codons under positive selection, and the NS2B:V103A mutation demonstrate ongoing viral adaptation. These findings highlight the need for continued genomic surveillance and targeted vector studies to guide WNV control strategies in Kenya and the wider region.

**Importance:** West Nile virus (WNV) remains a globally important arbovirus, yet genomic and experimental data from under-sampled regions such as coastal East Africa are limited. This study provides the first integrated molecular and genotypic characterization of WNV circulating along the Kenyan coast, revealing the co-detection of Lineage 1a and the first identification of Lineage 2 in this region. By combining field surveillance, whole-genome sequencing, evolutionary analyses, and lineage-specific replication assays across multiple vertebrate and mosquito cell lines, we demonstrate clear genetic and biological differences with implications for transmission and adaptation. Importantly, the detection of WNV in a male *Aedes aegypti* mosquito and recovery of full genomes offers compelling evidence of transovarial transmission, a mechanism that may support viral maintenance independent of vertebrate hosts. These findings expand current knowledge of WNV ecology in Africa and underscore the need for continued genomic surveillance to detect emerging variants and inform public health strategies.

## Introduction

West Nile virus (WNV) is a mosquito-borne virus belonging to the genus Flavivirus within the family Flaviviridae (1). It is a member of the Japanese encephalitis (JE) serocomplex (2). WNV is considered endemo-epidemic in many parts of the world, including Africa, Europe, the Middle East, Asia, and the Americas (3). It was first described in 1937 in Omogo, located in the West Nile district of northern Uganda, during a Yellow Fever surveillance campaign (4). Its continual geographic expansion poses a growing threat to the public (5). The primary vectors of WNV are mosquitoes, predominantly from the Culex genera (6). Although WNV has also been detected in other arthropods such as ticks and sandflies, their role in transmission remains unclear. A wide range of vertebrate hosts, especially birds from the order Passeriformes, can be infected. However, only a subset of these birds act as competent reservoirs, capable of sustaining and amplifying the virus (7). Mosquitoes acquire infection by feeding on infectious birds and, after a temperature-dependent extrinsic incubation period of 2 to 14 days, become infectious and can transmit the virus through subsequent blood meals (8). In this transmission cycle, mammals, particularly humans and horses, are incidental, dead-end hosts (9).

The WNV genome is approximately 11 kb in length and consists of a single-stranded, spherical, positive-sense RNA (10). It encodes a polyprotein that is cleaved into three structural (C, prM/M, and E) and seven non-structural (NS1, NS2A, NS2B, NS3, NS4A, NS4B, and NS5) proteins. It is genetically diverse and currently classified into at least nine lineages, with Lineages 1 (L1) and 2 (L2) being the most epidemiologically significant (5). Lineage 7 (L7) has been reclassified as a distinct flavivirus, Koutango virus (KOUTV), with a putative Lineage 8 also proposed (11). WNV Lineage 1 (L1) is the most widespread and genetically diverse, associated with outbreaks in humans, birds, and horses across continents. L1 is subdivided into clades 1a, 1b, and 1c. Clade 1a is globally distributed and includes strains from Africa, Europe, Asia, the Middle East, and the Americas (12). It was responsible for the 1999 outbreak in New York City, which marked WNV’s introduction into the Western Hemisphere (2). In Europe, clade 1a continues to cause recurrent outbreaks, with ongoing circulation and evolution in regions such as Italy and Spain. African strains of clade 1a have been detected in Senegal and Kenya, indicating persistent endemic transmission (13,14). Clade 1b, also known as Kunjin virus, is endemic to Australia and associated with milder disease, while clade 1c includes strains primarily found in India, contributing further to the virus’s genetic diversity (15). Lineage 2 (L2), once thought to be restricted to sub-Saharan Africa and associated with mild disease, has emerged as a significant cause of neuroinvasive disease in Europe (16). L2 is genetically distinct from L1 and is divided into clades 2a and 2b, with clade 2a further split into clusters A and B. Cluster B was linked to a major 2010 outbreak in Greece with a 17% case fatality rate, while cluster A emerged in the Netherlands in 2020, causing milder disease (12). The basis for differences in virulence between these clusters remains unclear.

WNV infection is predominantly asymptomatic. Around 20% of infected individuals may develop West Nile fever (WNF), characterized by influenza-like symptoms, and less than 1% develop West Nile neuroinvasive disease (WNND), which may include encephalitis, meningitis, acute flaccid paralysis, or death. Symptom severity depends on both host and viral factors, including strain virulence (17). In horses, symptomatic cases may develop severe neurological disease with a ∼33% mortality rate (2). Among birds, corvids and raptors are particularly susceptible, often dying from neurological complications. The virus replicates at the inoculation site, spreads to lymph nodes and the bloodstream, and can cross the blood-brain barrier, likely through inflammatory pathways involving Toll-like receptor stimulation and tumour necrosis factor-α–mediated increases in vascular permeability. WNV directly infects neurons, especially in the deep nuclei and gray matter of the brain, brainstem, and spinal cord (10).

WNV is endemic throughout Africa, where it circulates primarily among mosquitoes and avian hosts (18). Multiple lineages have been detected across the continent. Lineage 1 has been reported in Algeria, Egypt, Côte d’Ivoire, Kenya, Morocco, Senegal, and South Africa, while Lineage 2 has been implicated in severe human disease in Uganda, Tanzania, and South Africa (19). The Koutango lineage (WN-KOUTV), originally a separate flavivirus, has been identified in Senegal, the Central African Republic (11), and more recently in Kenya, isolated from sandflies in Baringo and Isiolo counties suggesting potential alternative transmission cycles (14). Evidence show that WNV may be maintained through vertical mechanisms such as transovarial transmission, where the virus is passed from an infected female mosquito to its offspring (20). Vertical transmission of WNV provide a mechanism for persistence during inter-epidemic periods. Major African outbreaks include the 1974 and 1980s epidemics in South Africa (21) and repeated equine and human outbreaks in North Africa between 1996 and 2012 (7). In Kenya, West Nile virus (WNV) has been isolated from various mosquito species collected across diverse geographical regions. This includes *Culex univittatus* from the Rift Valley (22), *Aedes sudanensis* from the North Eastern region (23), and multiple *Culex* and *Aedes* species during the 2007 Rift Valley fever outbreak in the same region (24). Additionally, WNV was detected in a pool of *Rhipicephalus pulchellus* ticks (5). The virus has also been detected in avian hosts (18), and a confirmed human infection further underscores the potential of WNV to pose a public health threat in Kenya (25).

Despite the growing body of knowledge on WNV in Kenya, surveillance still lacks consistency, and the true burden of the virus is not fully understood. As such, continued surveillance is critical to understanding WNV transmission in Kenya, tracking potential evolutionary trends of the virus, and identifying areas of high risk for outbreaks. Integrated vector management strategies, including the use of insecticides, biological control methods, and environmental management practices, are crucial for reducing the spread of WNV and other mosquito-borne diseases. Moreover, enhanced surveillance efforts in both urban and rural areas will be essential to capture a comprehensive picture of WNV circulation in Kenya. In addition to the classical bird–mosquito cycle, alternative transmission mechanisms such as transovarial transmission in mosquitoes have been suggested, which may contribute to viral persistence during inter-epidemic periods but remain poorly studied in Africa. This study contributes to the understanding of WNV genetic diversity in mosquito populations in Kenya, focusing on four counties: Kwale, Kilifi, Mombasa, and Isiolo. These regions were selected due to their ecological variability and history of arbovirus outbreaks. The study employs entomological sampling, cell culture, and sequencing methods to detect and characterize circulating viruses.

## Materials and Methods

### Ethical Approval

Ethical approval was obtained from the Kenya Medical Research Institute (KEMRI) Scientific and Ethics Review Unit (SERU) under protocol number KEMRI/SERU/CVR/4702 and WRAIR# 3101. Permission to conduct the study was granted by the National Council for Science, Technology, and Innovation (NACOSTI).

### Study area

The study was carried out in four Kenyan counties, Kilifi County (Malindi) (3.2192°S, 40.1169°E), Kwale (4.1730°S, 39.4520°E), Mombasa (4.0435°S, 39.6682°E), and Isiolo county (0.3546°N, 37.5822°E) (Fig 1).

**Figure 1.**
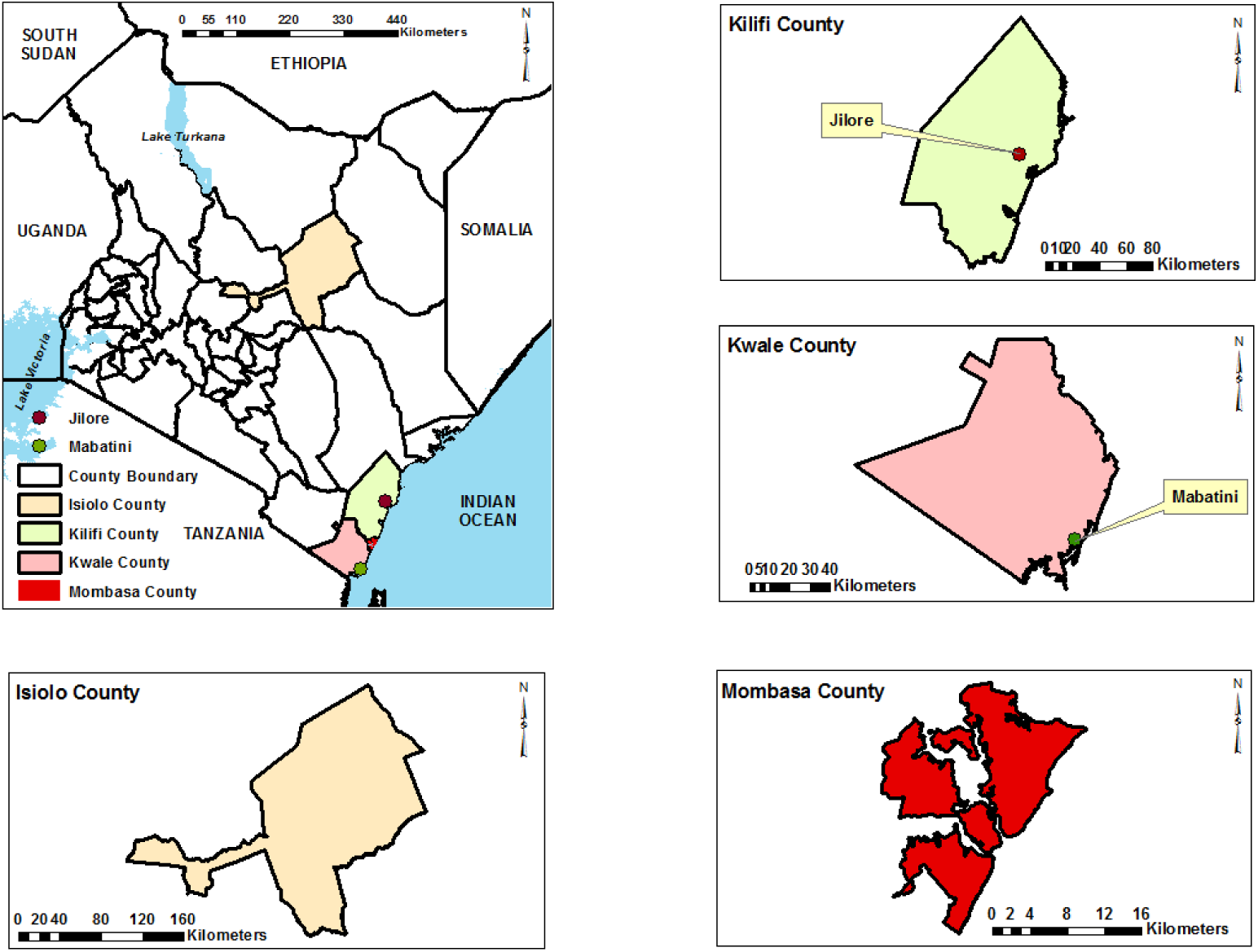
The overall map of coastal Kenya showing the site where sampling was conducted. Base maps, boundaries and shape files of Kenyan map and administrative boundaries of the County and Sub-county were derived from GADM data version 4.1 (https://gadm.org) and the maps were generated using ArcGIS Version 10.2.2 (http://desktop.arcgis.com/en/arcmap) advanced license) courtesy of Samuel Owaka.

### Entomological Investigation

Sampling of adult mosquitoes was conducted between 18^th^ June and 10^th^ December 2023. Mosquitoes were collected using CDC miniature light traps (Model 512, John Hock Co., Gainesville, Florida, USA). Traps baited with carbonated dry ice (CO2) were deployed overnight (6pm-6am) in favourable habitat, including dwelling quarters and animal sheds. The samples were linked to the sites by geo-coding using a GPS. The mosquitoes were immobilized by freezing at -20°C for 20mins and identified morphologically to species under a dissecting microscope using taxonomy keys, including Edwards (1941) (26), Harbach (1988)(27) and Jupp (1986) (28). The identified mosquitoes were pooled in groups of 1 to 25 samples based on species, sex and collection site. Mosquitoes were consequently preserved in liquid nitrogen, and transported to the laboratory at the Kenya Medical Research Institute in Kisumu, where they were stored at-80°C until further processing.

### Mosquito sample preparation

One thousand five hundred ninety-six (1,596) mosquito pools were homogenized using a Mini-Beadruptor-16 (Biospec, Bartlesville, OK, USA) in 1000 µL of homogenization media (minimum essential media supplemented with 15% fetal Bovine Serum (FBS) (Gibco by Life Technologies, Grand Island, NY, USA), 2% L-glutamine (Sigma, Aldrich)and 2% antibiotic/antimycotic (Gibco by Life Technologies, Grand Island, NY, USA.) with zirconium beads (2.0 mm diameter) for 40 seconds. Subsequently, the homogenates were centrifuged at 14,000 rpm for 15 minutes at 4°C using a benchtop centrifuge (Eppendorf, USA). The supernatant was collected and inoculated into Vero cells immediately.

### Viral Isolation

Virus isolation was performed in Vero (African green monkey kidney) cells (E6). Briefly, Vero cells were grown overnight at 37°C and 5% CO_2_ in minimum essential medium supplemented with 2% glutamine, 2% penicillin/streptomycin/amphotericin, 10% fetal bovine serum, and 7.5% NaHCO_3_ in 24-well plates (Corning, Incorporated). At 80% confluence, a 50 µL aliquot of the clarified supernatant from individual pools was inoculated into the wells. The plates were incubated for 1 hour in a humidified incubator at 37 °C and 5% CO_2_ with gentle rocking of the plates every 15 minutes for virus adsorption. Following incubation, 1 mL of maintenance medium, comprising minimum essential medium supplemented with 2% glutamine, 2% penicillin/streptomycin/amphotericin, 2% fetal bovine serum, and 7.5% NaHCO_3_, was added. The plates were then cultured at 37 °C and 5% CO_2_ and monitored daily for cytopathic effects (CPE) for 14 days. Cultures exhibiting CPE were harvested and further passaged by inoculating onto fresh monolayers of Vero cells (CCL 81™) in 25-cm^3^ cell culture flasks. After three successive passages, the supernatants of virus-infected Vero cell cultures exhibiting cytopathic effect of approximately 70% were harvested from the flasks for virus identification through next-generation sequencing.

### Library preparation and next generation sequencing

Viral particles from CPE-positive cultures were recovered through 0.22µm filters (Millipore, Merck). From an aliquot of 140 μl of the supernatant, viral RNA was extracted using the QIAamp Viral RNA Mini Kit (Qiagen, Hilden, Germany) and eluted in one step with 60 μl of elution buffer. Paired-end libraries for high-throughput sequencing were prepared using Illumina Ribo-Zero Plus rRNA Depletion (Illumina, USA). The libraries were prepared following the manufacturer’s recommended protocol. The final library was denatured with NaOH and diluted further to 10pM before loading onto the Illumina MiSeq sequencing machine. Sequencing was performed using MiSeq Reagent V3 (Illumina, USA), in a 300-paired end cycle sequencing format.

### Sequence analysis and virus identification

Initial analysis was performed using the CZ-ID platform, an integrated pipeline that offers quality control, de-hosting, duplicate removal, assembly, and viral identification capabilities. After this initial processing, the de-hosted sequence reads were retrieved for further analysis. To validate CZ-ID pipeline results, PrinseqLite v0.20.4 tool was used to filter low-quality reads and remove adapters on command line. De novo sequence assembly was conducted using MEGAHIT v1.2.9 (29) where West Nile virus contigs were recovered and compared to CZ-ID pipeline results. This was followed by mapping the reads back to the generated contigs using BWA to produce SAM files. The subsequent generation of BAM files, along with sorting, indexing, and filtering of contigs that did not meet the specified read mapping thresholds, was handled using Samtools v1.20. Only contigs with an average depth of coverage of ≥10 and a length of ≥500 bp were retained for further analysis. These contigs were first compared against a local version of NCBI viral database using Diamond v2.0.4. To ensure specificity, putative viral contigs were further compared to the entire non-redundant protein database (nr), to exclude any non-viral contigs. A stringent e-value threshold of 1e-5 was employed throughout the homology searches to minimize false-positive hits.

### Phylogenetic analysis of the identified RNA viruses

To describe the identified viruses in an evolutionary context, publicly available viruses belonging to these different groups, and more specifically those closely related to the viral strains obtained in the current study were downloaded and used as reference sequences in the reconstruction of phylogenetic trees. Closely related gene sequences were retrieved from NCBI viral database and used as reference sequences in reconstructing the phylogenetic relationship of the viral sequences. The combined set of sequences were aligned using MUSCLE software with default parameters (max iterations = 16) embedded in Molecular Evolutionary Genetics Analysis v.7.0 (MEGA7) (30) platform. The aligned sequences were edited using the Bioedit tool and maximum likelihood phylogenetic analysis carried out using IQ-TREE v1.6.12. The best model (GTR+G4 (General Time Reversible + Gamma)) and tree search was performed simultaneously based on 1000 bootstrap estimates and approximate likelihood ratio test (aLRT).

### Bayesian Evolutionary Analysis of WNV Lineages 1 and 2

To estimate the coalescent times of the most recent common ancestor (tMRCA) and the evolutionary relationships of West Nile Virus (WNV) lineages 1 and 2, molecular clock analysis was performed using the Bayesian Markov Chain Monte Carlo (MCMC) method implemented in BEAST v1.10.4 (Bayesian Evolutionary Analysis Sampling Trees). Representative complete genome sequences of WNV lineages 1 and 2, together with their country and year of isolation metadata were retrieved from the NCBI Viral Genome Database (https://www.ncbi.nlm.nih.gov/labs/virus/) and combined with sequences obtained in this study. The final dataset comprised 149 sequences for Lineage 1 and 122 sequences for Lineage 2, each with an alignment length of approximately 9,000 base pairs, aligned using MAFFT v7.

An uncorrelated lognormal relaxed molecular clock model was applied to accommodate rate variation among branches. The General Time Reversible (GTR) + Gamma + 4 substitution model and a Bayesian Skyline coalescent prior were used to estimate changes in effective population size through time.

Each MCMC analysis was run for 1 billion generations, sampling every 100,000th generation. Convergence and effective sampling were evaluated using Tracer v1.7.2, ensuring all parameters achieved effective sample sizes (ESS) > 200. The maximum clade credibility (MCC) tree was generated in TreeAnnotator v1.10.4, with the first 10% of samples discarded as burn-in. The resultant time-calibrated phylogenies were visualized using FigTree v1.4.4 (http://tree.bio.ed.ac.uk/software/figtree/), showing node ages and posterior probability support values at each node. The tMRCA estimates were represented as median years with 95% Highest Posterior Density (HPD) intervals.

### Growth Kinetics of WNV Lineage 1 and 2

The replication dynamics of West Nile Virus (WNV) Lineage 1 and Lineage 2 were evaluated in vertebrate (Vero Bikens, Vero CCL-81, and Vero E6) and mosquito-derived (C6/36) cell lines. Virus stocks, confirmed by whole-genome sequencing, were standardized to titres of 6.0 × 10⁶ PFU/mL for Lineage 1 and 3.7 × 10⁶ PFU/mL for Lineage 2. For vertebrate cells, monolayers were established in 24-well plates by seeding 2.0 × 10⁵ cells per well (Vero Bikens, Vero CCL-81, and Vero E6). Mosquito-derived C6/36 cells were seeded in T25 flasks at 2.8 × 10⁶ cells per flask to account for their growth requirements. Once confluent, cells were infected at a multiplicity of infection (MOI) of 0.1 with Lineage 1 and Lineage 2 virus, with gentle rocking every 15 minutes for 1 hour at the appropriate temperature, after which maintenance medium was added. Supernatants were harvested at 0, 12, 24, 36, 48, 60, 72, 84, 96, 108, and 120 hours post-infection (hpi) for Vero Bikens, Vero CCL-81, and E6 cells, while for C6/36 cells, supernatants were collected every 24 hours post-infection. Virus quantification was performed using standard plaque assays on Vero CCL-81 cells. Briefly, cells were seeded in 12-well plates and allowed to reach 100% confluency before infection with 100 µL of 10-fold serially diluted supernatants. After a 1-hour incubation at 37°C, cells were overlaid with a methylcellulose medium containing 12.5% agarose and 2% FBS and incubated at 37°C with 5% CO₂ for 4–6 days until plaques became visible. Cells were then fixed with 3.7% formaldehyde, stained with 0.5% crystal violet, and plaques were counted to calculate viral titres expressed as plaque-forming units per milliliter (PFU/mL). All experiments were performed in duplicate.

### Selection pressure analysis

To investigate the selection pressures acting on West Nile Virus (WNV) isolates recovered in this study, codon-based analysis was conducted using the Datamonkey Adaptive Evolution Server (http://www.datamonkey.org/). Full coding sequences were first aligned using MEGA7 software. Codon alignments were generated using HyPhy tools on the Datamonkey platform to preserve reading frame integrity. Three complementary methods were employed to identify codon sites under diversifying (positive) selection: FEL (Fixed Effects Likelihood) with a significance threshold of p ≤ 0.05 for detecting pervasive selection; MEME (Mixed Effects Model of Evolution) with p ≤ 0.01 for identifying episodic selection; and FUBAR (Fast Unconstrained Bayesian Approximation) with a posterior probability ≥ 0.99 for detecting pervasive selection using a Bayesian framework. Codon sites were considered positively selected if identified as significant by the three models. These sites were then mapped to the corresponding viral proteins to evaluate their potential role in viral evolution and adaptation.

### Amino Acid Variation

To investigate structural implications of amino acid variation in the West Nile Virus NS2B protein, the protein sequence obtained from our isolates was aligned with the reference sequence from NCBI RefSeq (Accession No: NC_009942.1) using MEGA7. A missense mutation was identified at codon position 103, resulting in a valine-to-alanine substitution (V103A). To model the 3D structure of the NS2B protein and assess the structural context of this mutation, AlphaFold v2.3.2 was used to predict the tertiary structure based on the amino acid sequence. The resulting structural model was then visualized and annotated using PyMOL v2.5.4.

### Species confirmation of mosquito pools

COI gene amplification and sequencing were conducted to confirm the morphological identification of mosquito species in each pool. DNA extraction was performed on the bulk pools of the crude homogenates using a QIAamp DNA extraction kit (Qiagen, Germany), according to the manufacturer’s instructions. Two extraction blanks were included during the extraction process and subsequently used during PCR and sequencing. The extracted DNA was quantified using a Qubit fluorometer 2.0. Amplification of the COI gene was then carried out on <1µg of extracted DNA, using the universal pair of primers for metazoan invertebrates LCO1490/HCO2198 (31). These primers amplify an approximately 710-bp region of the COI gene of arthropod vectors. COI amplicons were generated from a 25µL PCR containing 12.5 µl AmpliTaq Gold 360 master mix (Applied Biosystems, USA), 9.5µl DNase/RNase-free water, and 0.5µl each of the forward and reverse primers at 25µM. The PCR cycling conditions were set as follows; initial denaturation at 95°C for 10 min, 35 cycles of 95°C for 30 s, 49°C for 30 s, 72°C for 30 s, and a final extension of 72°C for 10 min. The COI amplicons were first purified using AMPure XP beads (Beckman Coulter, USA). The purified products were quantified using a Qubit dsDNA HS assay kit (Invitrogen, USA) with a Qubit fluorometer 2.0. Based on the concentration of the quantified products, the volume of PCR products that yielded 200 fmol was determined and used as starting material for MinION library preparation. Library preparation was carried out using a ligation sequencing kit (SQK-LSK114), following the manufacturers protocol with the exclusion of the DNA fragmentation step. Briefly, 200 fmol of the purified products were end repaired using a NEBNext Ultra II end repair and dA-tailing module (New England Biolabs [NEB], UK). The end-repaired DNA for each sample was individually barcoded using Native Barcoding Expansion 1-12 (EXP-NBD114), which was achieved with the use of NEB Blunt/TA ligase master mix (NEB, UK). An equal amount from each of the 200-fmol barcoded libraries was combined into a single pool, which was then purified with AMPure XP beads (Beckman Coulter, USA). Adapter ligation of the purified library was done with NEBNext quick ligation module (NEB, UK) and the libraries were further purified using AMPure XP beads, with a final wash of the beads being carried out using short fragment buffer (SFB) provided with the SQK-LSK114 kit. The final library was loaded onto the flow cell (FLO-MIN106D) and sequenced using the workflows provided in the MinKNOW software. Raw reads were processed in real-time using the EPI2ME platform (Run on cloud), which performed base-calling with Guppy (v4.2.2), quality filtering (minimum Q-score 7), and taxonomic classification using the ‘What’s in my Pot’ (WIMP) workflow. Taxonomic assignments were validated by BLAST searches against the NCBI nucleotide database and cross-checked with the Barcode of Life Data System (BOLD) for species confirmation. Only sequences with ≥98% identity and ≥90% query coverage were considered reliable identifications.

## Results

### Viral Isolation and Whole Genome sequencing

A total of 14,105 mosquitoes were grouped into 1,596 pools according to species and geographic origin. Ten (10) pools exhibited cytopathic effects (CPE) after three successive passages in Vero E6 cells. Infected cultures developed CPE by day 5 post-inoculation, characterized by cell rounding, detachment, and loss of the monolayer. Sequencing of the CPE-positive pools revealed West Nile Virus (WNV) in all ten isolates, with eight complete genomes and two partial genomes recovered. Genome typing identified WNV Lineage 1a in eight pools and Lineage 2 in two pools across the surveyed regions.

**Table 1:** Viruses identified in this study based on observed cytopathic effects (CPE) and sequence analysis. The identification was carried out using a homology search against reference databases, providing insights into the closest known virus.

| *CPE Positive*<br>*pools* | *Region* | *Species* | *Sex* | *WNV*<br>*Lineage* | *Length*<br>*(bp)* |
| --- | --- | --- | --- | --- | --- |
| MAL/S3/44066 | Kilifi | *Culex pipiens* | F | 1a | 11,137 |
| MAL/S9/44235 | Kilifi | *Anopheles funestus* | F | 1a | 6,958 |
| MBSA/S3/44491 | Mombasa | *Aedes aegypti* | F | 2 | 10,982 |
| MBSA/S2/44358 | Mombasa | *Aedes aegypti* | M | 1a | 10,989 |
| MBSA/S1/44288 | Mombasa | *Culex univittatus* | F | 2 | 2,394 |
| MBSA/S3/44496 | Mombasa | *Aedes aegypti* | F | 1a | 11,102 |
| MBSA/S2/44439 | Mombasa | *Eretmapodites*<br>*chrysogaster* | F | 1a | 10,989 |
| KWL/S5/44632 | Kwale | *Aedes aegypti* | F | 1a | 10,999 |
| KWL/S1/44904 | Kwale | *Aedes aegypti* | F | 1a | 10,981 |
| KWL/S1/44901 | Kwale | *Aedes aegypti* | F | 1a | 11,006 |

### Phylogenetic analysis

Phylogenetic analysis was performed to describe and infer lineages of the WNV isolated from the mosquito pools in this survey in relation to other strains isolated available in GenBank. Analysis showed that eight pools clustered within West Nile Virus (WNV) Lineage 1, Clade A, forming a branch with the 2012 Senegal strain. The remaining two isolates clustered within WNV Lineage 2, branching from the Ugandan strain.

**Figure 2:**
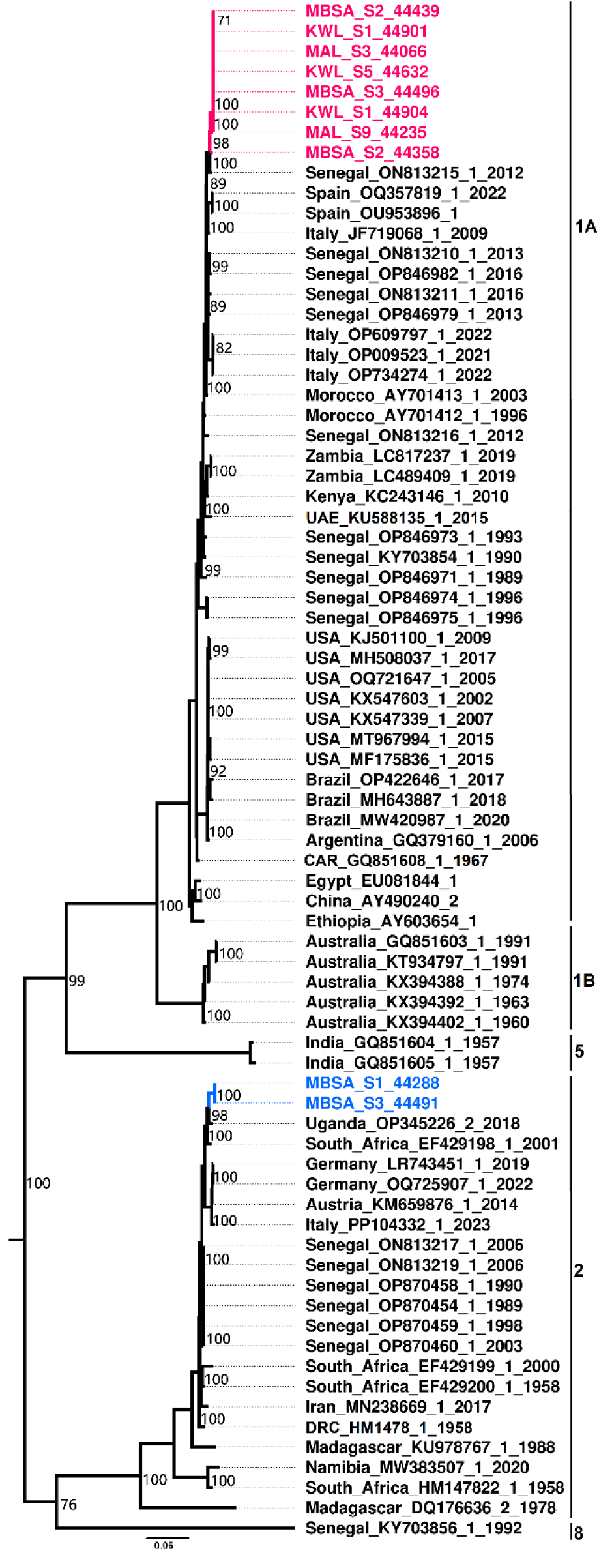
Maximum likelihood phylogenetic tree constructed using a nucleotide sequence data set including those obtained in this study (red and blue). Isolates highlighted in red cluster within Lineage 1a, while blue-highlighted isolates clustered within Lineage 2. Bootstrap support values from 1000 replicates are indicated on the branches.

### Bayesian Evolutionary Analysis of West Nile Virus Lineage 1

The time-calibrated phylogeny inferred using BEAST revealed that the eight (8) West Nile Virus (WNV) lineage 1 isolates from this study clustered within the lineage 1a clade. These isolates formed a well-supported monophyletic cluster branching from Italian strains sampled around 2010.64 (95% HPD: 10.35). One (1) isolate showed a distinct evolutionary placement, diverging more recently around 2020.66 (95% HPD: 3.92) suggesting an independent introduction event or recent local evolution within the region.

**Figure 3.**
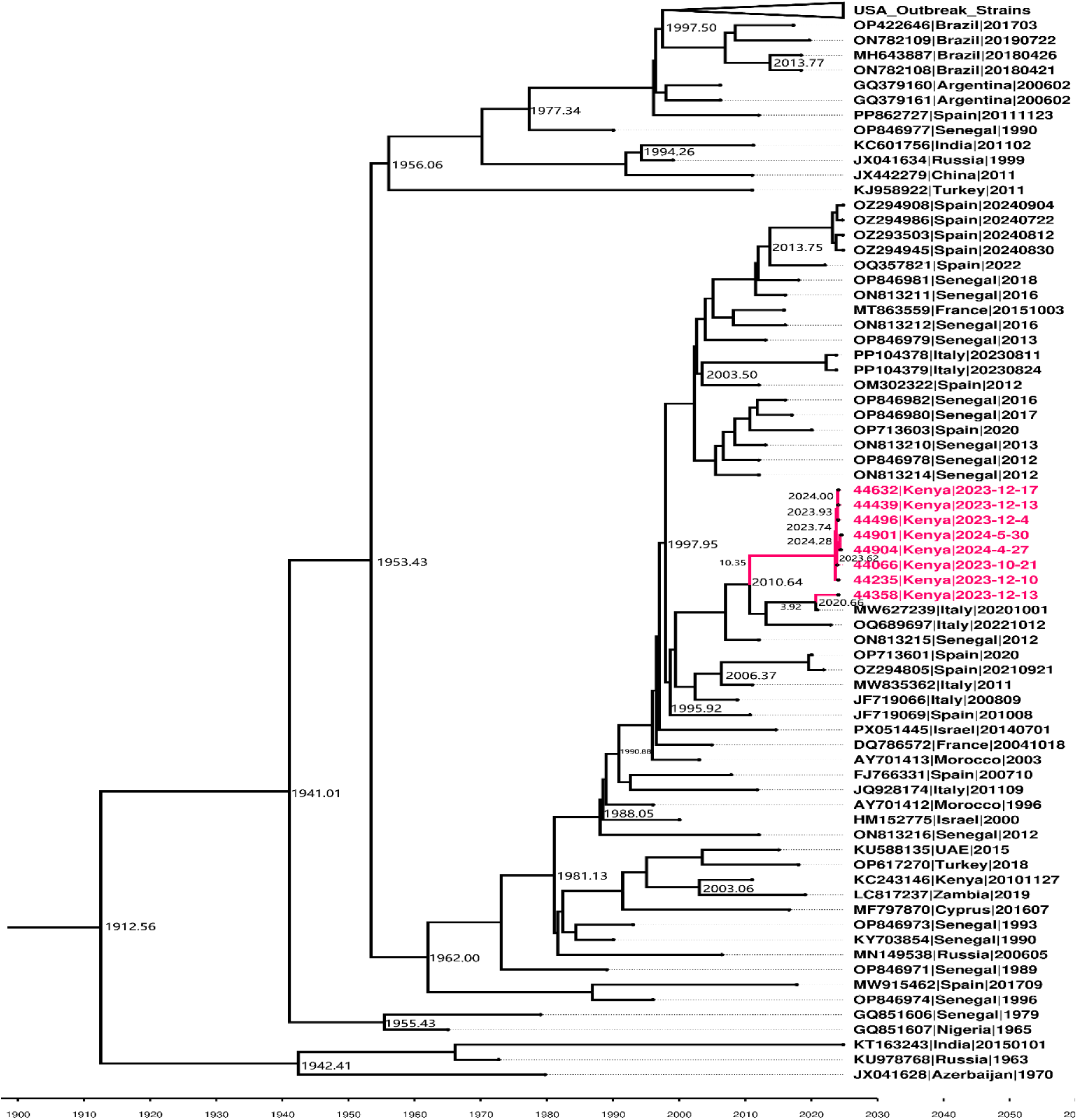
Time-Calibrated Maximum Clade Credibility (MCC) tree of West Nile Virus (WNV) Lineage 1 inferred using BEAST v1.10.4. Kenyan isolates from this study are highlighted in red and cluster within the lineage 1a clade, branching from Italian strains sampled around 2010.64, while one isolate diverged around 2020.66. Node bars represent the 95% Highest Posterior Density (HPD) intervals for the estimated time to the most recent common ancestor (tMRCA).

### Bayesian Evolutionary Analysis of West Nile Virus Lineage 2

The Bayesian time-calibrated phylogeny revealed that the two West Nile Virus (WNV) lineage 2 isolates from this study clustered within the South African lineage 2 clade. The isolates shared a common ancestor with South African strains that circulated around 1994.38, indicating a long-term evolutionary link with southern African WNV populations. The two Kenyan isolates formed a distinct, well-supported subclade that diverged from each other around 2023.56 (95% HPD: 0.89) suggesting recent diversification or local evolution.

**Figure 4.**
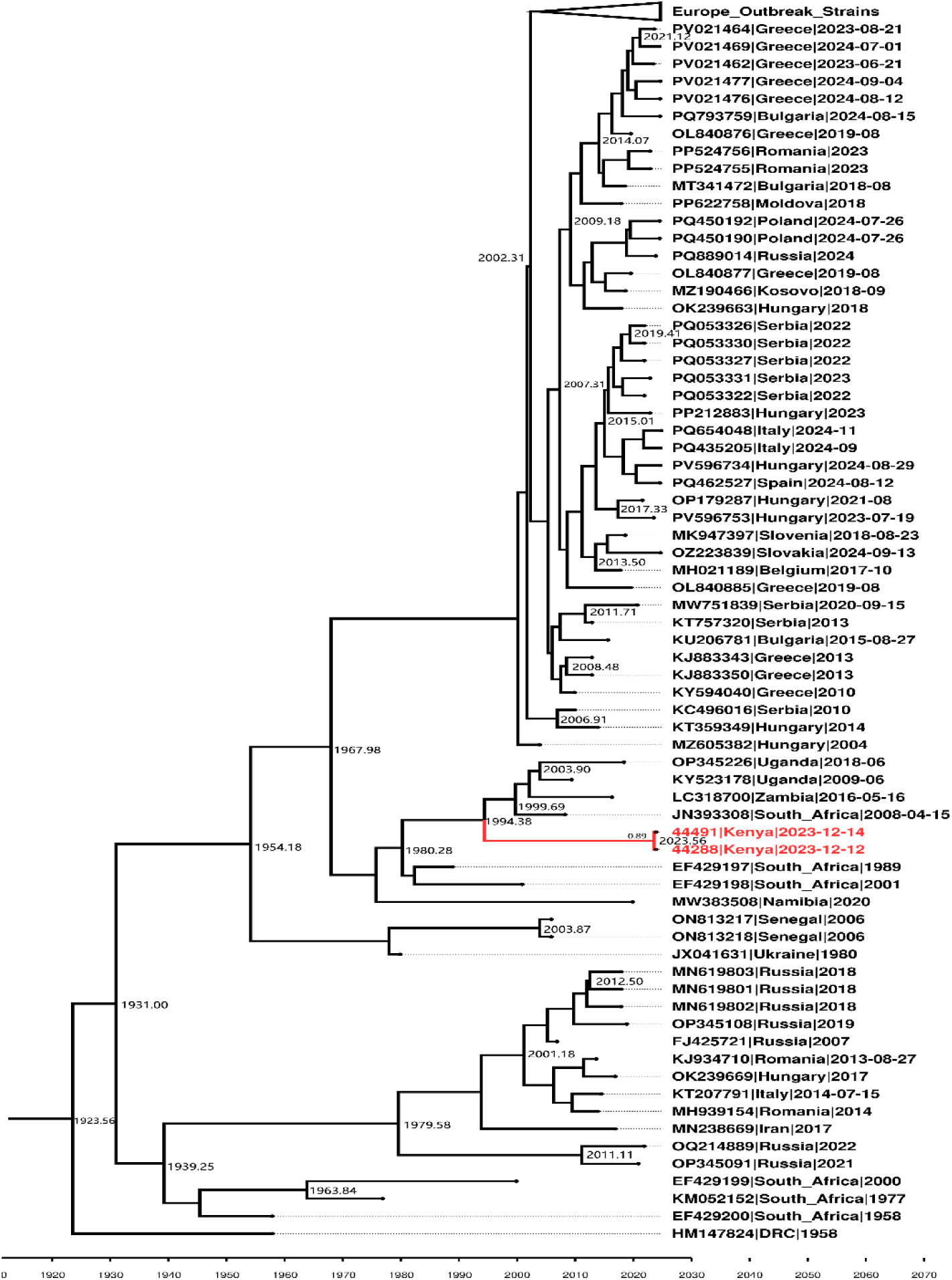
Time-calibrated maximum clade credibility (MCC) tree of West Nile Virus (WNV) lineage 2 inferred using BEAST v1.10.4. The two Kenyan isolates from this study are highlighted blue and clustered within the South African lineage 2 clade, diverging from South African strains around 1994.38. The two isolates subsequently branched from each other around 2023.56, indicating recent diversification.

### Growth Kinetics of WNV Lineage 1 and 2

Distinct replication profiles were observed between WNV lineages and cell types. Growth kinetics assays were performed in replicates, with MBSA/S3/44496 selected as the representative Lineage 1 isolate and MBSA/S3/44491 selected as the representative Lineage 2 isolate (no specific criteria applied during isolate selection). Overall, Lineage 1 achieved higher peak titres than Lineage 2 in Vero Bikens, Vero E6 and C6/36 cells, while Lineage 2 replicated slightly better in Vero CCL-81. In mosquito-derived C6/36 cells, both lineages reached rapid early replication; however, Lineage 1 maintained higher titres throughout the infection period. Biological replicates showed high reproducibility, with peak titres varying by approximately 0.5 log units.

**Figure 5.**
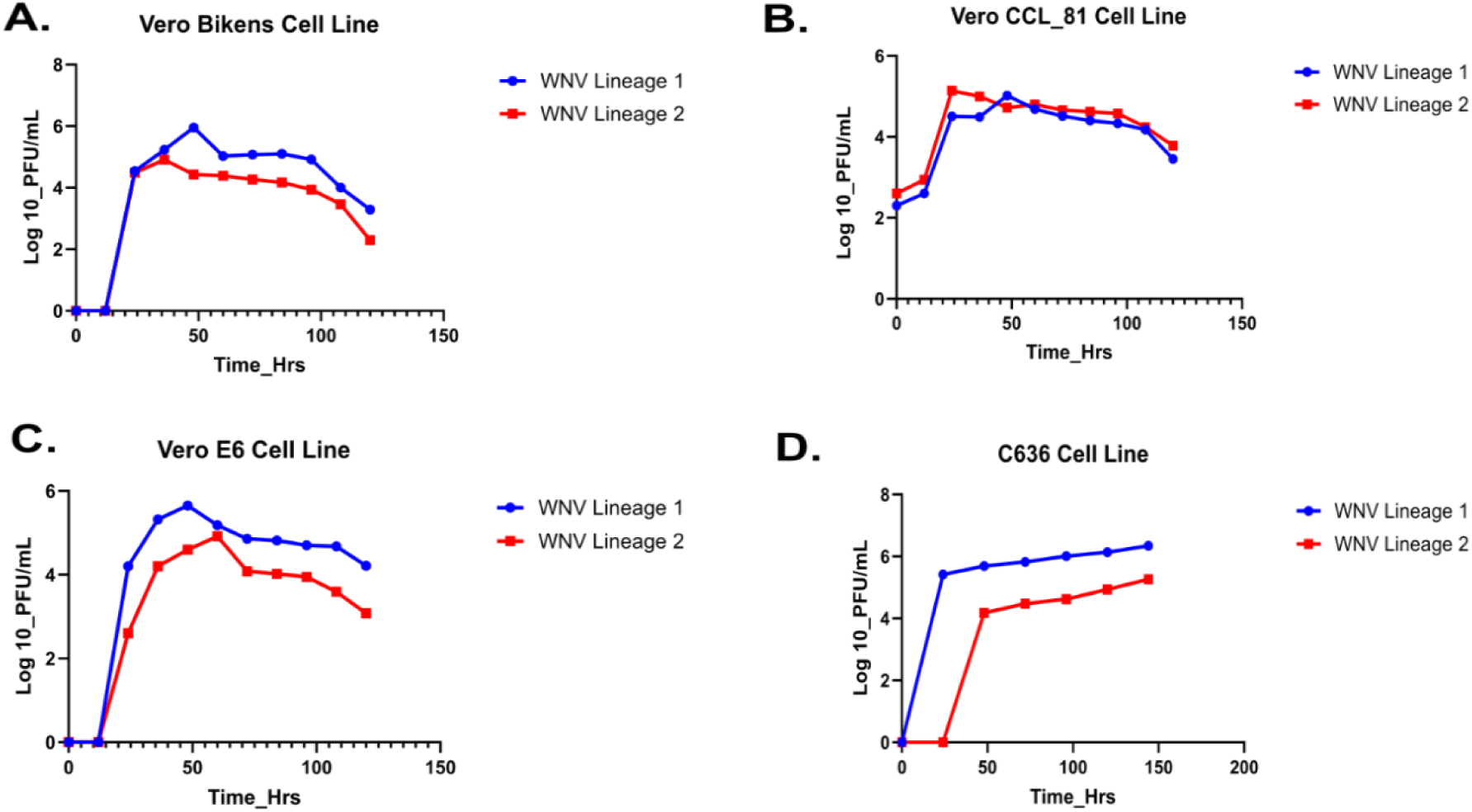
Replication kinetics of WNV Lineage 1 and Lineage 2 in four different cell lines. Viral growth curves showing infectious virus titres (PFU/mL) over time post-infection in (A) Bikens, (B) Vero CCL-81, (C) Vero E6, and (D) C6/36 cells. Overall, Lineage 1 achieved higher peak titres than Lineage 2 in Bikens, Vero E6 and C6/36 cells, while Lineage 2 replicated slightly better in Vero CCL-81. C6/36 cells supported the highest replication overall, with peak titres observed later in infection compared to mammalian cell lines.

**Table 2.** Peak titres and time for West Nile Virus Lineage 1 and Lineage 2 isolates as shown in different mammalian and insect cell lines.

| Cell Line | Lineage 1<br>Peak titre<br>(PFU/ml) | Peak time<br>(hpi) | Lineage 2<br>Peak titre<br>(PFU/ml) | Peak time<br>(hpi) |
| --- | --- | --- | --- | --- |
| **Bikens** | $9.0 \times 10^5$ | 48 hpi | $8.2 \times 10^4$ | 36 hpi |
| **CCL-81** | $1.0 \times 10^5$ | 48 hpi | $1.4 \times 10^5$ | 24 hpi |
| **Vero E6** | $4.5 \times 10^5$ | 48 hpi | $8.2 \times 10^4$ | 60 hpi |
| **C6/36** | $2.2 \times 10^6$ | 144 hpi | $1.8 \times 10^5$ | 144 hpi |

### Selection Pressure

The Datamonkey platform consistently identified diversifying (positive) selection on six codon sites. These sites included codons 119 and 188 in the NS2A gene, codon 103 in NS2B, and codons 44,177, and 422 in the RNA-dependent RNA polymerase (RdRp) region. Notably, our isolates harbored the NS2B:V103A mutation, which was also found to be under significant positive selection.

**Figure 6.**
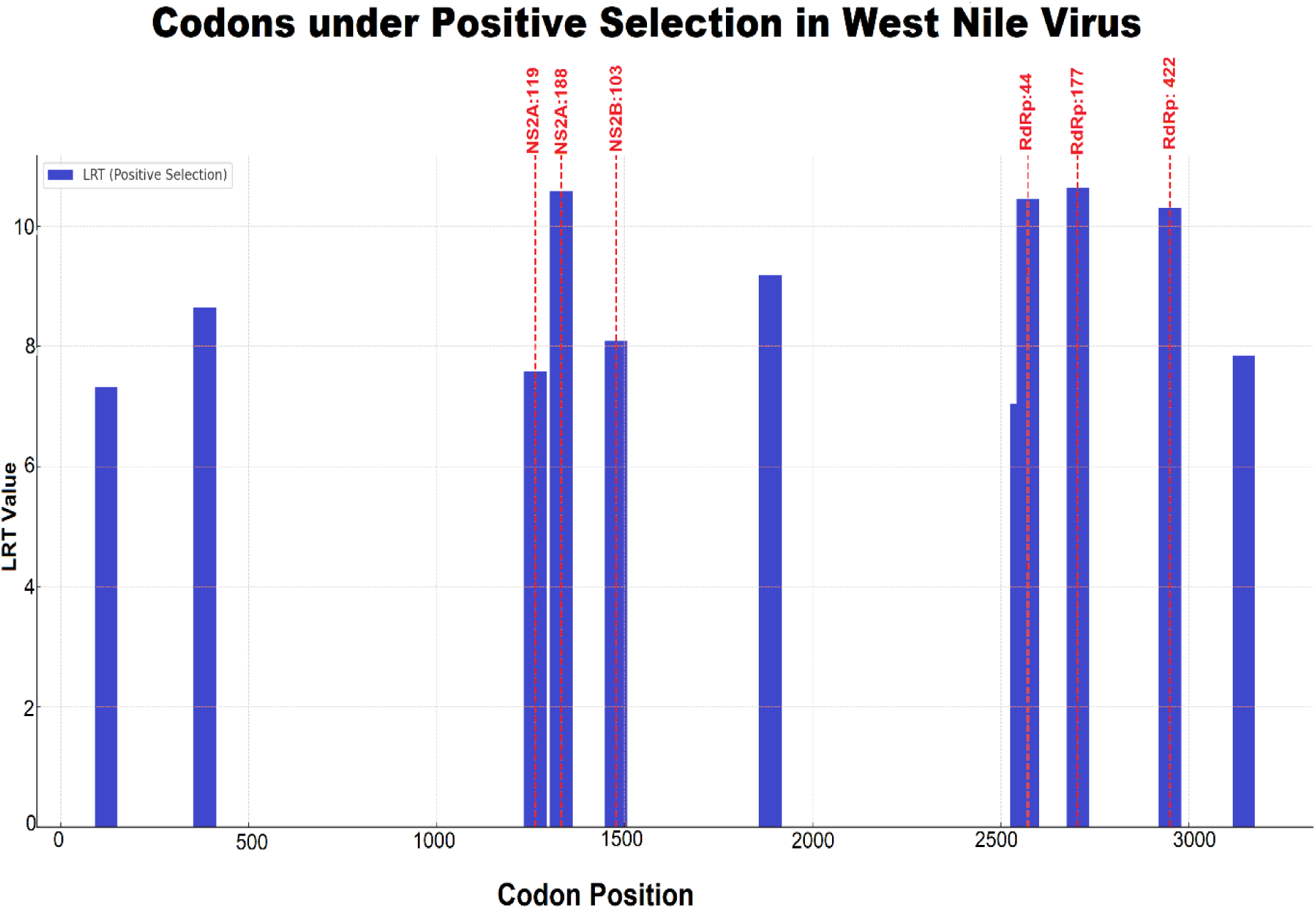
Codon sites under positive selection in West Nile Virus genomes identified using Datamonkey. Positively selected sites were detected at codons 119 and 188 in NS2A, codon 103 in NS2B, and codons 44, 177, and 422 within the RNA-dependent RNA polymerase (RdRp) region.

### Amino Acid Variation

The NS2B protein sequence obtained from our isolates was compared to the reference sequence available in NCBI RefSeq (Accession No: NC_009942.1). Sequence alignment revealed a missense mutation at position 103, where valine (V) was substituted with alanine (A) (V103A). The resulting 3D structure confirmed the presence of the V103A substitution, as shown in Figure 7 below.

**Fig 7:**
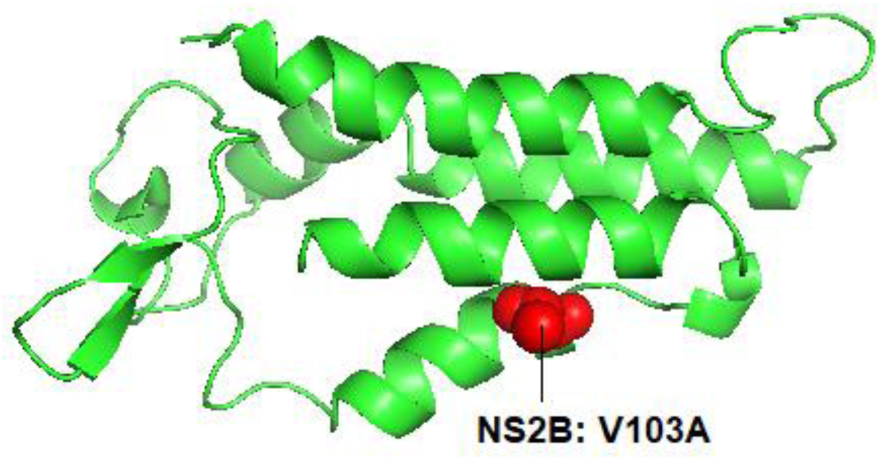
AlphaFold-predicted three-dimensional structure of the West Nile Virus NS2B protein highlighting the V103A amino acid substitution. The identified positively selected site at codon 103 is shown in pink, illustrating the spatial positioning of the mutation within the NS2B structural domain.

### Metabarcoding

Sequencing was successful for all the 10 pools. Approximately 873,541 raw sequence reads, with the read distribution among the 6 regions as given below, were sequenced (Table 3). The average quality scores for the sequenced reads ranged from 17 to 20 and the median lengths of the reads for the pools ranging from 600 to 702 base pairs.

**Table 3:** Comparison of Morphological and COI Molecular Identities of Sampled mosquitoes.

| CPE Positive pools | Morphological Identity | Molecular Identity | Pool size |
| --- | --- | --- | --- |
| MAL/S3/44066 | *Culex pipiens* | *Culex pipiens* | 15 |
| MAL/S9/44235 | *Anopheles funestus* | *Anopheles funestus* | 17 |
| MBSA/S3/44491 | *Aedes aegypti* | *Aedes aegypti* | 11 |
| MBSA/S2/44358 | *Aedes aegypti* | *Aedes aegypti* | 7 |
| MBSA/S1/44288 | *Culex univittatus* | *Culex univittatus* | 13 |
| MBSA/S3/44496 | *Aedes aegypti* | *Aedes aegypti* | 6 |
| MBSA/S2/44439 | *Eretmapodites chrysogaster* | *Eretmapodites chrysogaster* | 4 |
| KWL/S5/44632 | *Aedes aegypti* | *Aedes aegypti* | 1 |
| KWL/S1/44904 | *Aedes aegypti* | *Aedes aegypti* | 1 |
| KWL/S1/44901 | *Aedes aegypti* | *Aedes aegypti* | 3 |

## Discussion

West Nile virus (WNV) remains a significant emerging arboviral threat with public health implications globally and in sub-Saharan Africa. This study contributes to ongoing surveillance efforts by providing molecular and phylogenetic evidence of WNV presence and diversity in four ecologically diverse counties of Kenya, namely: Kwale, Kilifi, Mombasa and Isiolo. Of the 1,596 mosquito pools screened, ten (10) showed cytopathic effects (CPE) on Vero E6 cells and yielded high-quality WNV sequences upon analysis, confirming the reliability of combining cell culture with molecular detection to identify infectious virus.

Several species of mosquitoes were found to harbor WNV, including *Culex pipiens*, *Culex univittatus*, *Anopheles funestus*, *Aedes aegypti*, and *Eretmapodites chrysogaster*. Notably, *Culex* species are recognized as the primary enzootic vectors of WNV, especially *Culex pipiens* and *Culex univittatus*, which have previously been implicated in both zoonotic and epidemic cycles in Africa and beyond (10). The presence of WNV in *Aedes aegypti*, *Anopheles funestus*, and *Eretmapodites* species, traditionally not considered primary vectors, raises questions about the vector competence and ecological dynamics that may support spillover in diverse environments. While incidental, these findings underscore the need for further studies to elucidate the potential role of these species in WNV ecology. Additionally, detection of WNV Lineage 1 in a male *Aedes aegypti* specimen with a full genomic sequence strongly implies a non-blood acquisition pathway and therefore supports the possibility of vertical transmission within Kenyan *Aedes* populations. This is an important inference for understanding persistence and overwintering potential. Vertical transmission of WNV has previously been demonstrated in mosquitoes (22) supporting the biological plausibility of this mechanism in our context.

Phylogenetic analysis revealed that the isolates clustered into two major phylogenetic groups: Lineage 1 (n=8) and Lineage 2 (n=2). The presence of Lineage 2 in *Culex univittatus* and *Aedes aegypti* in Mombasa represents the first detection of L2 in this region of Kenya. Previous surveillance historically indicated predominance of Lineage 1 in eastern Africa(32), whereas Lineage 2 circulation has been more commonly associated with southern Africa and Europe (33). Our findings therefore expand the ecological range of Lineage 2 in East Africa and strongly support contemporary dissemination, ecological expansion, or renewed introduction events.

Bayesian evolutionary analyses further identified distinct temporal diversification among isolates. Lineage 1 viruses clustered within the 1a clade, branching from strains previously seen in Italy around 2010.64. Conversely, the two Lineage 2 viruses clustered within the South African clade, sharing a recent common ancestor around 1994.38 and diverging between themselves around 2023.56. This supports a model of multiple introductions or long-term cryptic maintenance followed by recent micro-evolutionary divergence, consistent with recent re-emergence patterns reported in other African contexts (2). The global molecular clock for WNV further supports rapid evolutionary turnover potential, emphasizing that Kenyan lineages must be monitored longitudinally to distinguish one-time incursions from persistent establishment. The temporal diversification observed underscores the dynamic nature of WNV transmission and the need for continuous genomic surveillance to detect emerging strains, track lineage movement, and inform vector control and public health strategies aimed at mitigating future outbreaks.

Our study highlights distinct lineage and cell-specific replication kinetics of West Nile Virus (WNV) Lineage 1 and Lineage 2 in mammalian (Vero Bikens, Vero CCL-81, Vero E6) and mosquito (C6/36) cell lines. These findings underscore that viral replication is strongly influenced not only by lineage differences but also by the cellular environment. The faster replication of Lineage 1 in Vero Bikens and Vero E6 aligns with previous reports that L1 strains often achieve higher titres and earlier peaks in vertebrate cells compared to L2 (34,35). Interestingly, Vero CCL-81 cells displayed an opposite trend, with Lineage 2 peaking as early as 24 hpi, while Lineage 1 peaked later. Such discrepancies suggest that cell line–specific factors, including receptor availability, endocytic pathways, or innate antiviral responses, may differentially shape WNV replication, as observed in other flavivirus studies (33,34). In mosquito-derived C6/36 cells, both lineages established productive infection, though Lineage 1 reached higher titres. This is consistent with the intrinsic permissiveness of C6/36 cells, linked to their impaired RNA interference pathway (36). Nevertheless, lineage-dependent differences in replication efficiency in mosquito cells have also been noted (37), indicating that vector competence may vary depending on the specific lineage–vector pairing. Collectively, our results reinforce that the replication dynamics of WNV are shaped by an interplay between viral genetic background and host cell environment. The observed differences between Lineage 1 and Lineage 2 across cell lines may have implications for transmission dynamics, virulence, and adaptation during enzootic and epidemic cycles.

The NS2B:V103A substitution, observed in all isolates, was particularly noteworthy and under significant positive selection. NS2B acts as a cofactor for the NS3 protease, and substitutions in this region could affect viral replication or interaction with host immune responses (38,39). Structural modeling confirmed the location of the V103A mutation but did not reveal gross conformational disruption, suggesting functional rather than structural significance. The recurrence of this mutation across isolates may indicate adaptive evolution in response to host or vector pressures.

In addition to WNV, the Kenyan coastal region is increasingly recognized as a hotspot for the co-circulation of multiple flaviviruses, including dengue virus and a diverse assemblage of insect-specific viruses (ISVs) (40). Co-circulation of flaviviruses is virologically significant because concurrent infections in shared vector species can drive viral interference, modulate superinfection exclusion, and influence the RNA interference (RNAi)–mediated antiviral landscape of mosquitoes (34). ISVs, in particular, have been shown to alter replication kinetics of medically important flaviviruses by competing for cellular machinery or priming antiviral pathways, which may affect vector competence for WNV (41). Additionally, overlapping transmission cycles increase the likelihood of heterologous selection pressures, which could shape adaptive mutations such as those observed in WNV NS2B. The presence of WNV in an ecological context rich in other flaviviruses therefore highlights a complex multispecies viral network that may influence viral fitness, transmissibility, and evolutionary trajectories, underscoring the need for integrated genomic surveillance of co-circulating arboviruses.

## Conclusion

This study provides strong evidence of active co-circulation of two pathogenic WNV lineages in Kenyan mosquito populations, identifies the first detection of Lineage 2 in coastal Kenya, demonstrates lineage-dependent replication phenotypes in vitro, and reveals ongoing adaptive evolution in key non-structural proteins. The recovery of a full genomic sequence of Lineage 1 from a mal*e Aedes aegypti* further supports the possibility of transovarial transmission within Kenyan populations, strengthening the inference that non-blood acquisition pathways contribute to local maintenance in the region. These findings emphasize the need for continued integrated genomic, vector ecological, and evolutionary surveillance to anticipate, detect, and mitigate future WNV emergence and outbreak risk in Kenya and across the broader Eastern African region.

## Recommendations

- Prioritize vector competence experiments using Kenyan *Aedes aegypti* and *Culex* populations for both WNV Lineage 1 and Lineage 2 to quantify lineage-specific transmission efficiency, midgut escape, and salivary gland infection profiles.
- Integrate mosquito sex-stratified screening in national arbovirus surveillance pipelines to strengthen detection of possible vertical transmission signals and improve understanding of non-blood dependent maintenance pathways.

## Ethics approval and consent to participate

Ethical approval was obtained from the Kenya Medical Research Institute (KEMRI) Scientific and Ethics Review Unit (SERU) under protocol number KEMRI/SERU/CCR/4702 and WRAIR# 3101. Permission to conduct the study was granted by the National Council for Science, Technology, and Innovation (NACOSTI).

## Acknowledgements

We thank, Victor Ofula, Dr. Samson Konongoi, Francis Mulwa, Dr. Edith Chepkorir, Simon Muhoro and Joseph Katur for their expert contribution in cell culture and data analysis.

## Disclaimer

This Material has been reviewed by the Walter Reed Army Institute of Research. There is no objection to its presentation and/or publication. The opinions or assertions contained herein are the private views of the author, and are not to be construed as official, or as reflecting true views of the Department of the Army or the Department of Defense.

## Availability of data and materials

The sequences of the viruses identified in this study have been submitted to GenBank with accession numbers: PX619830, PX619831, PX619832, PX619833, PX619834, PX619835, PX619836, PX619837, PX619838, PX619839.

